# Synaptotagmin 1 oligomers clamp and regulate different modes of neurotransmitter release

**DOI:** 10.1101/594051

**Authors:** Erica Tagliatti, Oscar D. Bello, Philipe R. F. Mendonça, Dimitrios Kotzadimitriou, Elizabeth Nicholson, Jeff Coleman, Yulia Timofeeva, James E. Rothman, Shyam S. Krishnakumar, Kirill E. Volynski

## Abstract

Synaptotagmin1 (Syt1) synchronises neurotransmitter release to action potentials acting as the fast Ca^2+^ release sensor and as the inhibitor (clamp) of spontaneous and delayed asynchronous release. Whilst the Syt1 Ca^2+^ activation mechanism has been well characterised, how Syt1 clamps transmitter release remains enigmatic. Here we show that C2B domain-dependent oligomerisation provides the molecular basis for the Syt1 clamping function. This follows from the investigation of a designed mutation (F349A), which selectively destabilises Syt1 oligomerisation. Using combination of fluorescence imaging and electrophysiology in neocortical synapses we show that Syt1^F349A^ is more efficient than wild type Syt1 (Syt1^WT^) in triggering synchronous transmitter release but fails to clamp spontaneous and Synaptotagmin7 (Syt7)-mediated asynchronous release components both in rescue (Syt1^−/−^ knock-out background) and dominant-interference (Syt1^+/+^ background) conditions. Thus we conclude that Ca^2+^-sensitive Syt1 oligomers, acting as an exocytosis clamp, are critical for maintaining the balance among the different modes of neurotransmitter release.

## Introduction

Tightly regulated synaptic release of neurotransmitters forms the basis of neuronal communication in the brain. Synaptotagmin1 (Syt1) plays a key role in this process, both as the major Ca^2+^ sensor for fast synchronous action potential-evoked transmitter release and as an inhibitor of spontaneous and delayed asynchronous release. Syt1 is a synaptic vesicle-associated protein containing a large cytosolic part composed of two tandemly arranged Ca^2+^-binding C2 domains (C2A and C2B). The C2B domain also binds SNARE (soluble N-ethylmaleimide sensitive factor attachment protein receptor) complexes and acidic lipids on the presynaptic membrane, and thus supports the generation and maintenance of the readily releasable pool (RRP) of vesicles docked at synaptic active zone^1–3^. Action potential-evoked depolarisation triggers opening of presynaptic voltage-gated Ca^2+^ channels resulting in a transient and spatially restricted increase of [Ca^2+^] at the vesicular release sites. Upon Ca^2+^ binding the adjacent aliphatic loops on Syt1 C2 domains insert into the presynaptic membrane and this triggers rapid (sub-millisecond time-scale) fusion of RRP vesicles^4, 5^.

In physiological conditions Syt1 also acts as a suppressor (or ‘clamp’) of spontaneous transmitter release (that occurs in the absence of neuronal spiking) and of the delayed lasting increase in vesicular exocytosis that follows an action potential (asynchronous release). Indeed, at many synapses genetic deletion of Syt1 not only abolishes the fast synchronous release component but also leads to a several-fold enhancement of spontaneous and asynchronous release^6–8^. Similarly, deletion of the SNARE-binding presynaptic protein complexin partially abrogates the clamping phenotype, suggesting that a synergistic action of both Syt1 and complexin is required for the fusion clamp^9, 10^. Experimental evidence argues that the Ca^2+^-dependent triggering and clamping actions of Syt1 are distinct in molecular terms^2, 11, 12^. However, the precise molecular mechanism of the Syt1-mediated clamp remains the subject of debate^13, 14^.

We have recently demonstrated that isolated cytoplasmic portions of Syt1 readily polymerise into ring-like oligomers *in vitro*^15, 16^. Syt1 oligomerisation is triggered by binding of PIP2 (phosphatidylinositol 4,5-bisphosphate) or ATP to a conserved polybasic motif in the C2B domain, which has been implicated in vesicle docking both *in vitro* and *in* vivo^16^. The Syt1 oligomers formed on membrane surfaces are sensitive to Ca^2+^ and readily dissociate as the Ca^2+^-binding loops located at the dimer interface (Fig. 1a) reorient and insert into the membrane^15, 16^.

**Figure 1.**
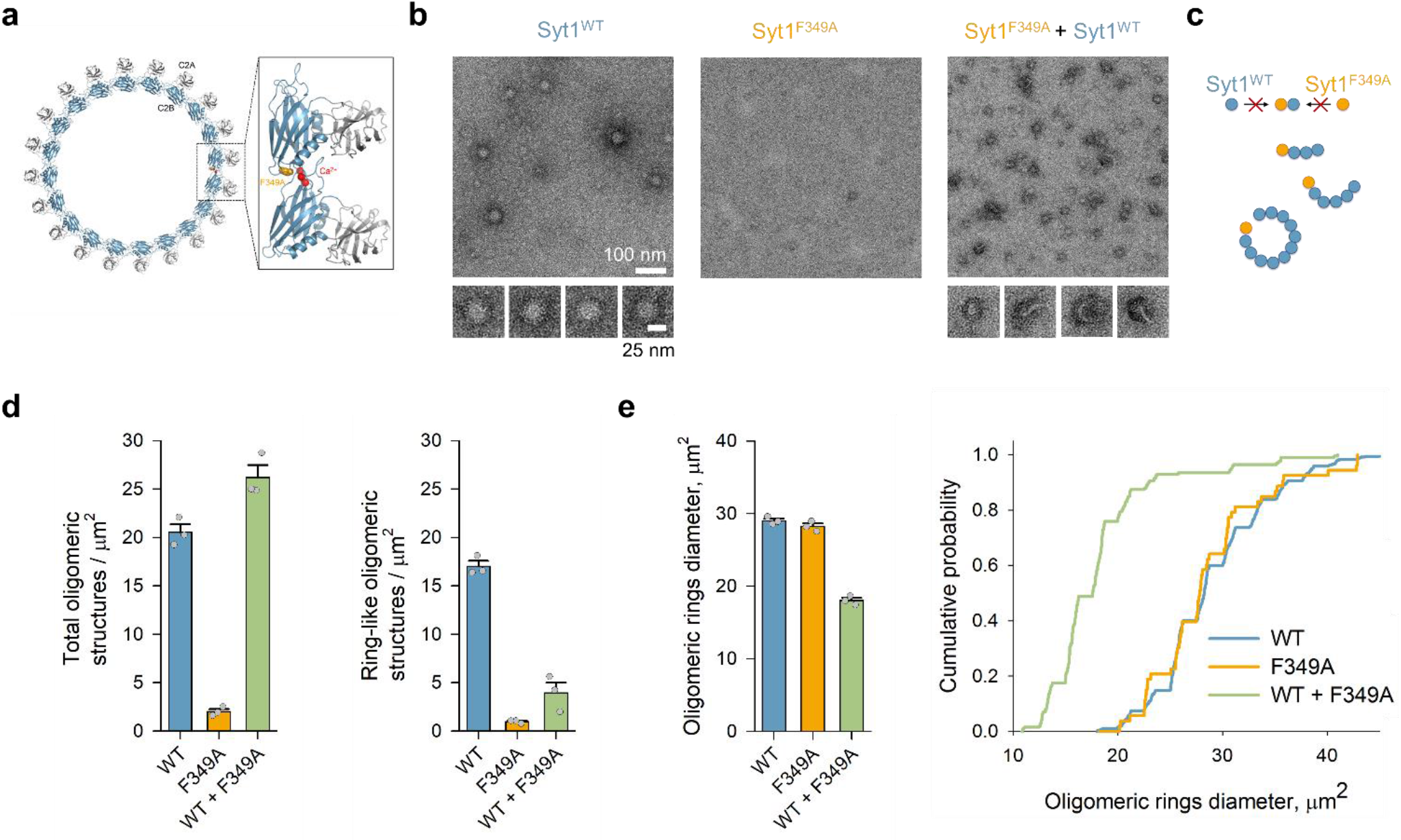
Dominant effect of F349A mutation on Syt1 oligomerisation. **(a)** Reconstruction of the Syt1 ring-like oligomers shows that the interaction of C2B domains (blue) drives the oligomerisation, with the Ca^2+^-binding loops (red) locating to the dimer interface. The F349A mutation designed to disrupt oligomerisation^17^ is shown in orange. **(b, c)** Negative-stain EM analysis shows that Syt1^F349A^ by itself does not form oligomeric structures, and when mixed with Syt1^WT^ (1:1 molar ratio) dominantly interferes with the formation of Syt1 oligomeric rings (b), likely by acting as a chain-terminator or the polymerisation reaction (c). **(d)** Syt1^F349A^, when mixed with Syt1^WT^ (1:1 molar ratio), does not change the total number of oligomeric structures (left), but critically lowers the number of ring-like oligomeric structures (right). **(e)** The resultant oligomeric ring-like structures were also smaller in size (average outer diameter ∼20 nm) as compared to ring-oligomers observed with Syt1^WT^ or Syt1^F349A^ alone (∼30 nm outer diameter). Bar graphs data are mean ± s.e.m. from 3 independent experiments. Cumulative plot shown in (d) is from 365 (Syt1^WT^), 53 (Syt1^F349A^) and 200 (Syt1^WT^ + Syt1^F349A^) ring-like particles pooled together from all experiments. The detailed statistical analysis is shown in Supplementary Table 1.

To assess, whether Syt1 oligomers have a physiological role, we designed a targeted mutation (F349A) that specifically destabilises Syt1 oligomerisation *in vitro,* without affecting other critical molecular properties, namely Ca^2+^ and SNARE binding and membrane interaction^17^. Previously we found that rescue overexpression of Syt1^F349A^ in non-neuronal neuroendocrine PC12 cells increased constitutive exocytosis, effectively occluding Ca^2+^-stimulated vesicular release^17^. We also observed that the F349A mutation abrogates the ability of Syt1 to clamp Ca^2+^-independent fusion in a reconstituted single vesicle fusion assay^18^. Together, these findings prompted the hypothesis that Syt1 oligomerisation could play a key role in mediating its clamping function in neuronal synapses.

Here we tested this hypothesis by systematically investigating the effects of the oligomer-destabilising F349A mutation on different modes of transmitter release in neocortical synapses in culture. We find that Syt1^F349A^ is more efficient than wild type Syt1 (Syt1^WT^) in rescuing synchronous transmitter release from knock-out (Syt1^−/−^) neurons, but fails to restore the clamp on asynchronous and spontaneous release. Furthermore, consistent with its dominant-interfering phenotype *in vitro,* overexpression of Syt1^F349A^ in Syt1^+/+^ neurons potentiates Syt1-mediated synchronous, Syt7-mediated asynchronous and action potential-independent spontaneous release components. Our data argue that Ca^2+^ sensitive Syt1 oligomers act as an exocytosis clamp and thus play the major role in coordinating different modes of transmitter release in central synapses.

## Results

### Dominant effect of F349A mutation on Syt1 oligomerisation

Based on the reconstruction of the Syt1 ring-like oligomer from EM density map, we previously designed the F349A mutation that selectively disrupts Syt1 ability to form oligomeric structures^17^. Prior to testing the effect of this mutation on neurotransmitter release, we further characterised the structural properties of Syt1^F349A^ protein. Negative stain electron microscopy analysis confirmed that the Syt1^F349A^ protein lacks the inherent ability to oligomerise and further revealed that it dominantly interferes with oligomerisation of Syt1^WT^ molecules (Fig. 1b). This is consistent with the model that Syt1^F349A^ acts as a chain terminator of the polymerisation reaction (Fig. 1c). With Syt1^WT^ alone, we predominantly observed stable ∼30 nm diameter size ring-like oligomers. In contrast, in a 1:1 Syt1^WT^/Syt1^F349A^ mixture, we observed a majority (>80%) of short, linear or curved polymers, with a minority (<20%) of smaller (∼ 20 nm diameter) ring-like structures, even though the total number of oligomers structures were comparable under both conditions (Fig. 1d, e).

### Disruption of Syt1 oligomerisation increases release in response to single action potentials

To assess the impact of interfering with Syt1 oligomerisation on vesicle exocytosis, we compared the functional effects of Syt1^F349A^ and Syt1^WT^ expression in either Syt1^−/−^ deficient or Syt1^+/+^ neurons (rescue and dominant interference experiments, respectively) (Fig. 2 and Supplementary Figs. 1 and 2). We monitored exocytosis by imaging the pH-sensitive fluorescent vesicular recycling probe synaptophysin-pHluorin (sypHy)^19^ (Fig. 2). We first recorded the increase in sypHy fluorescence in response to a burst of 20 action potentials at 100 Hz (ΔF_20APs_), which identified active synaptic boutons and gave an estimate of the relative size of the RRP at different synapses^20^. We also monitored the increase of sypHy fluorescence after cessation of the stimulation and used this as a measure of the delayed asynchronous release component (ΔF_Asynch_). Finally, we measured the presynaptic sypHy response (averaged over 10 trials) to single action potential stimulation (ΔF_1AP_), and estimated the average release probability of individual RRP vesicles as *p_v_* = ΔF_1AP_/ΔF_20APs_ (Fig. 2a)^20^.

**Figure 2.**
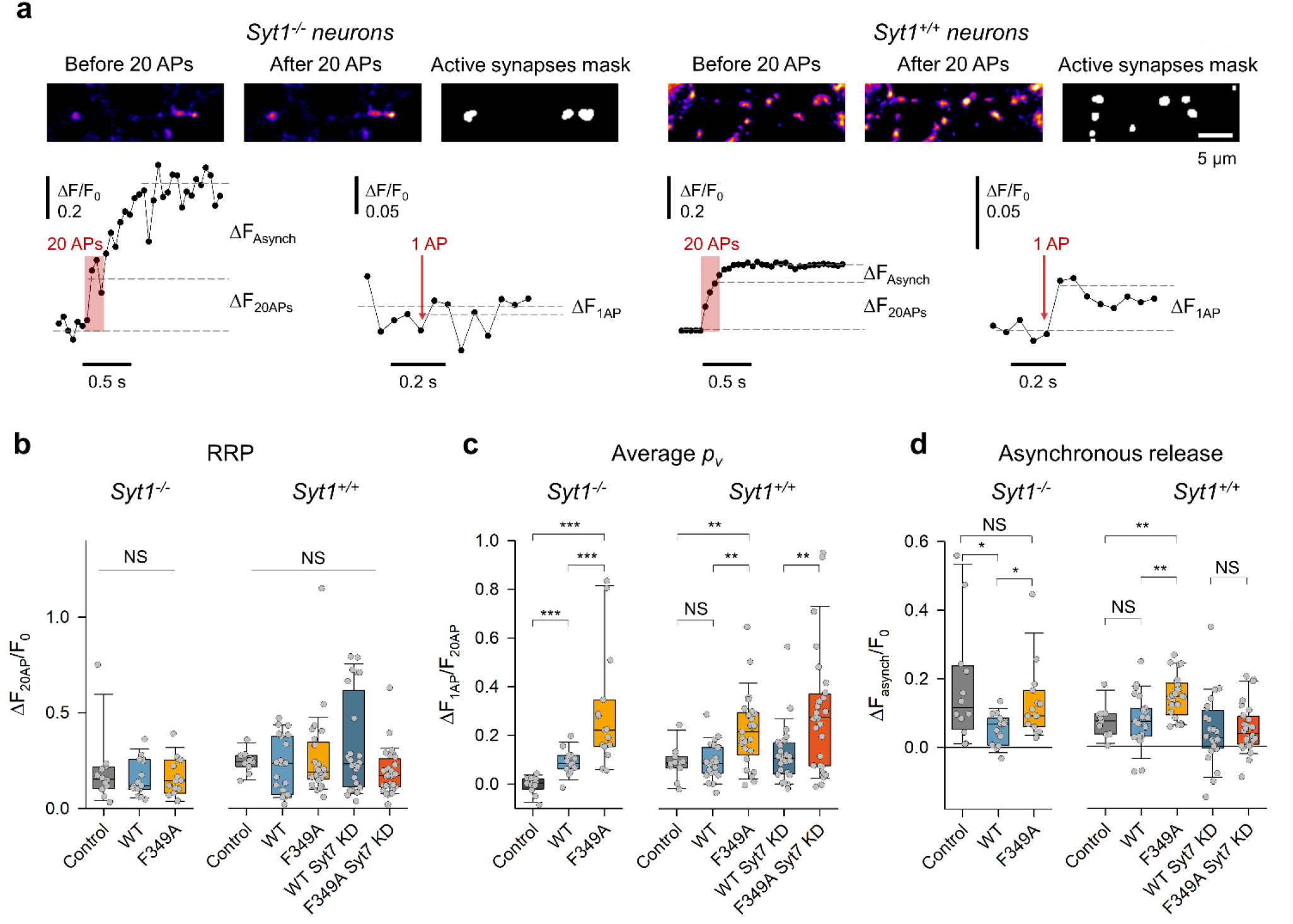
Disruption of Syt1 oligomerisation increases release in response to single action potentials and abolishes clamping of Syt7-mediated asynchronous release. **(a)** Representative sypHy fluorescence imaging experiments in Syt1^−/−^ and Syt1^+/+^ neurons designed to determine the relative RRP size (ΔF_20APs_), average release probability of RRP vesicles in response to a single action potential (AP) (*p_v_* = ΔF_1AP_/ΔF_20APs_)^20^, and the asynchronous release component (ΔF_Asynch_). **(b-d)** Summary box-and-dot plots showing that overexpression of Syt1^F349A^ does not change the RRP size **(b)** but increases *p_v_* **(c)** and the Syt7-mediated asynchronous release component **(d)** in both rescue (Syt1^−/−^) and dominant interference (Syt1^+/+^) experiments. * p < 0.05, ** p < 0.01, *** p < 0.001, NS p > 0.2, ANOVA on ranks **(b)**, and Mann–Whitney U test **(c, d)**. The detailed statistical analysis including the numbers of independent experiments is reported in Supplementary Table 1.

ΔF_20APs_ responses were similar in Syt1^WT^ and Syt1^F349A^ expressing synapses on both Syt1^−/−^ and Syt1^+/+^ backgrounds, suggesting that oligomerisation of Syt1 is not essential for generation and maintenance of the RRP (Fig. 2b). In line with the previously reported inhibition of synchronous transmitter release triggered by single spikes in Syt1 deficient synapses^21, 22^ we could not detect sypHy fluorescence increase in response to single action potential stimulation in Syt1^−/−^ neurons (Fig. 2a, c). However, Syt1^WT^ fully rescued the release triggered by single spikes to the same level as in wild type Syt1^+/+^ neurons (Fig. 2c). Syt1^F349A^ was approximately two-fold more efficient in rescuing release than the WT construct. Similarly, expression of Syt1^F349A^ in Syt1^+/+^ neurons resulted in a comparable increase of *p_v_* (Fig. 2c). The observed gain-of-function phenotype of Syt1^F349A^ suggests that Syt1 oligomerisation negatively regulates vesicular release triggered by single spikes.

### Disruption of Syt1 oligomerisation abolishes clamping of Syt7-mediated asynchronous release

ΔF_Asynch_ was approximately two-fold higher in Syt1^−/−^ than in Syt1^+/+^ synapses (Fig. 2d), in line with the reported potentiation of asynchronous transmitter release in Syt1^−/−^ neurons^8^. Expression of Syt1^WT^ on a Syt1^−/−^ background restored the clamping of the asynchronous release component, but Syt1^F349A^ failed to do so. Correspondingly, overexpression of Syt1^F349A^ on a Syt1^+/+^ background resulted in approximately two-fold potentiation of ΔF_Asynch_ in comparison to the untransduced control and Syt1^WT^ overexpression (Fig. 2d). Considering that asynchronous release in central synapses is largely mediated by the high-affinity Ca^2+^ sensor Syt7^8, 23^ we tested the effect of F349A mutation in Syt7 knock-down (KD) neurons. The increase of ΔF_Asynch_ induced by the F349A mutation was occluded by siRNA-induced knock-down (KD) of Syt7 (Fig. 2d and Supplementary Fig. 3). This result indicates that Syt1 oligomers clamp the Syt7-mediated asynchronous release component.

### Differential effects of Syt1^F349A^ on synchronous and asynchronous release components during repetitive stimulation

The majority of the release triggered by single spikes in glutamatergic neocortical synapses is tightly synchronised to action potentials^24^. However, due to the limited temporal resolution of sypHy imaging experiments (∼ 50 to 100 milliseconds) it was initially unclear to what extent the increase in *p_v_* observed in Syt1^F349A^ expressing neurons (Fig. 2c) was due to an increase of synchronous or asynchronous release components. To address this question, we used the recently generated fluorescence glutamate sensor SF-iGluSnFR.A184V (iGluSnFR) optimised for fast optical imaging of vesicular exocytosis^25^.

We first established a whole-cell current-clamp recording in an iGluSnFR expressing neuron and then imaged action potential-evoked release (triggered by somatic current injections) in tens of individual boutons supplied by its axon at a rate of ∼ 6 milliseconds per frame (Fig. 3a and related Supplementary Movie 1). We used 51 × 5 Hz stimulation to identify all boutons (including those with low release probability) and to measure the changes in synchronous and asynchronous release during repetitive spiking. To determine the precise timing of release events we applied a deconvolution procedure that took into account the temporal profile of uniquantal release events (see Methods). The responses at each release site that occurred within three frames (∼18 ms) after a given spike were classified as synchronous events (blue circles in Fig. 3a), and the rest as asynchronous events (red circles / arrows in Fig. 3a). The amplitude of the signals after deconvolution also allowed us to determine the quantal content of each release event.

**Figure 3.**
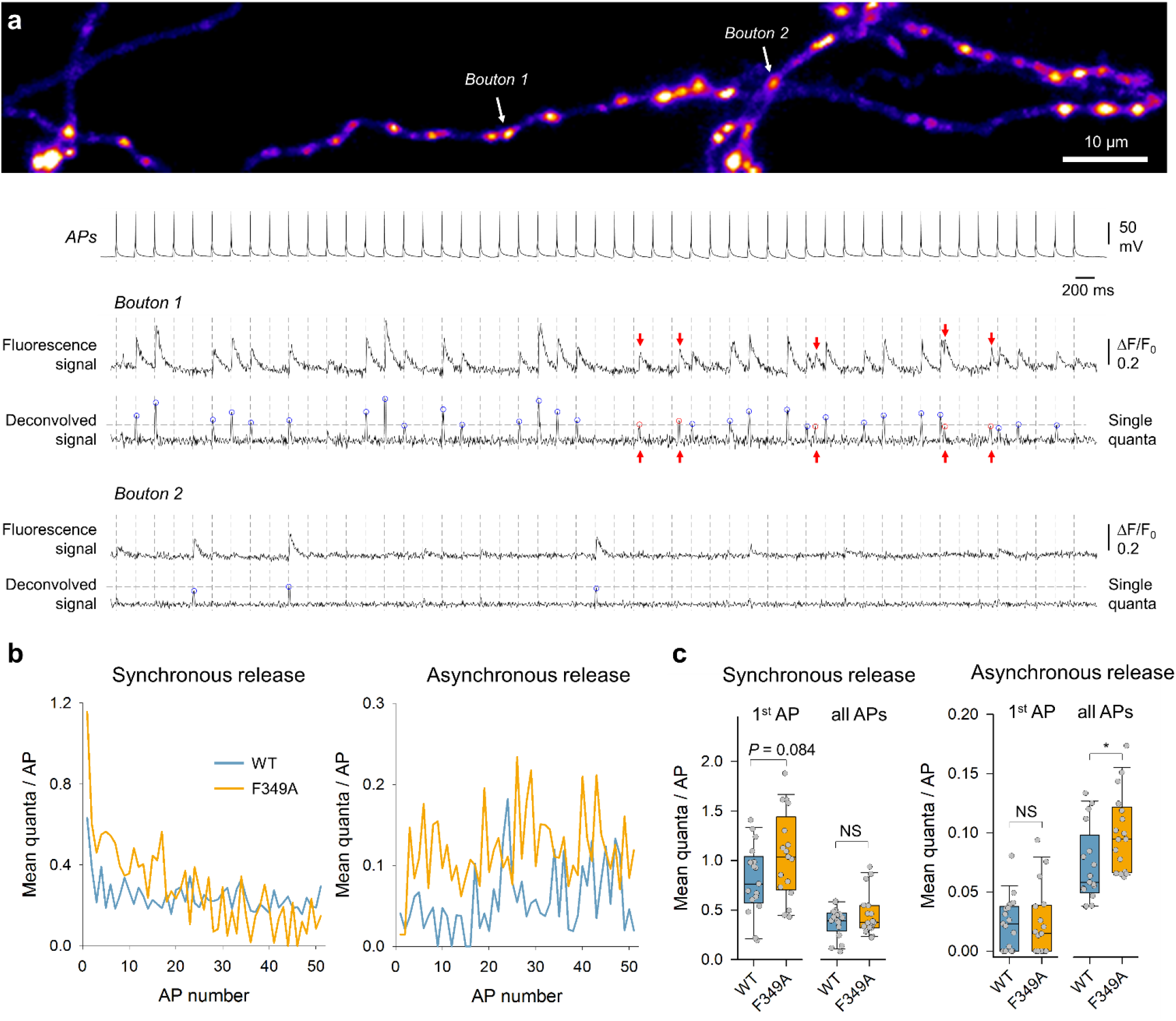
Differential effects of Syt1^F349A^ on synchronous and asynchronous release components during repetitive stimulation. **(a)** Representative iGluSnFR imaging experiment in control Syt1^+/+^ neurons, designed to detect quantal synchronous and asynchronous glutamate release events in individual synaptic boutons. Top, heat-map image revealing locations of glutamate release sites across the axonal arbour of a pyramidal neuron determined with 51 × 5 Hz stimulation (see Supplementary Movie 1). Traces, somatic action potentials and iGluSnFR fluorescence responses from two representative boutons with high and low release probabilities. Corresponding deconvolved signals were used to estimate the quantal size (horizontal grey lines) and the timing of release events (synchronous – blue circles, asynchronous – red circles / arrows). **(b)** Mean quantal responses for synchronous and asynchronous release at each spike in two representative experiments recorded in Syt1^+/+^ neurons overexpressing either Syt1^WT^ (n = 56 boutons) or Syt1^F349A^ (n = 62 boutons). **(c)** Summary plots showing the gain-of-function phenotype of the F349A mutation on synchronous release at the first spike and on asynchronous release triggered by the repetitive stimulation. Data points are mean values across all boutons (∼ 30 – 100 range) in each recorded cell. * p < 0.05, NS p > 0.7 Mann–Whitney U test. The detailed statistical analysis including the numbers of independent experiments is shown in Supplementary Table 1.

With the first action potential of the train, the synchronous component accounted for over 95% of all release in both Syt1^WT^- and Syt1^F349A^-overexpressing Syt1^+/+^ neurons. In line with the sypHy experiments, the F349A mutation caused an increase in the average number of quanta released with the first spike (Fig. 3b, c), albeit to a somewhat smaller extent than the increase in *p_v_* estimated using sypHy imaging (Fig. 2c). During repetitive stimulation the synchronous release showed prominent short-term depression (characteristic of neocortical glutamatergic synapses in culture^24^), effectively occluding the difference between Syt1^WT^- and Syt1^F349A^-expressing synapses.

In contrast, the asynchronous release component progressively increased during the stimulation, with approximately 1.6-fold stronger potentiation in Syt1^F349A^- than Syt1^WT^-expressing neurons (Fig. 3b, c). Taken together, the sypHy and iGluSnFR experiments show that destabilisation of Syt1 oligomers by the F349A mutation potentiates both Syt1-mediated synchronous and Syt7-mediated asynchronous release. Thus, we conclude that Syt1 oligomers act as negative regulators of both components of action potential-evoked release.

### Syt1 oligomerisation is required for clamping spontaneous release

Lastly, we determined the effect of the F349A mutation on spontaneous release in the absence of action potentials. We recorded miniature excitatory postsynaptic currents (mEPSCs) in the wholecell voltage-clamp configuration in the presence of the Na^+^ channel blocker tetrodotoxin (Fig. 4). The mEPSC frequency in Syt1^−/−^ neurons was several-fold higher than that in Syt1^+/+^ neurons, consistent with the proposed role of Syt1 in clamping spontaneous release^6, 7^. The clamp on spontaneous release was rescued in Syt1^−/−^ neurons by expression of Syt1^WT^ but not Syt1^F349A^. Correspondingly, overexpression of Syt1^F349A^ on a Syt1^+/+^ background resulted in approximately two-fold potentiation of spontaneous release (Fig. 4b). These results argue for the critical role of Syt1 oligomerisation in clamping spontaneous neurotransmitter release. The average amplitudes of mEPSCs were similar across all the conditions tested (Fig. 4c), ruling out any possible contribution or event misdetection to the observed differences in mEPSC frequencies.

**Figure 4.**
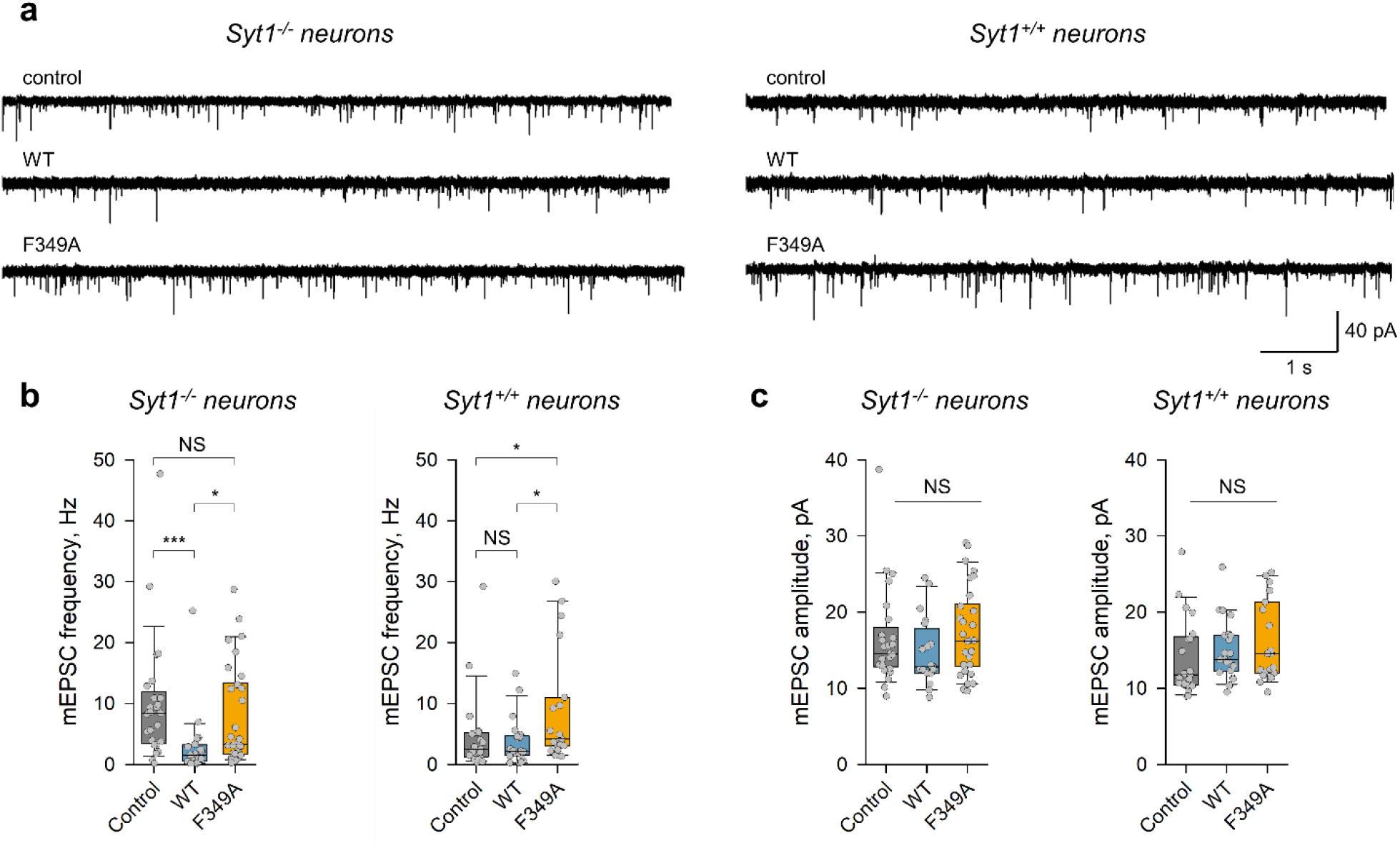
Syt1 oligomerisation is required for clamping spontaneous release. **(a)** Representative mEPSC traces recorded in untransduced Syt1^−/−^ or Syt1^+/+^ neurons (control) and in Syt1^−/−^ or Syt1^+/+^ neurons overexpressing either Syt1^WT^ or Syt1^F349A^. **(b, c)** Quantification of the effects of Syt1^WT^ or Syt1^F349A^ overexpression on mEPSC frequency **(b)** and amplitude **(c)**. Syt1^F349A^ fails to rescue the Syt1-mediated clamp of spontaneous release in Syt1^−/−^ neurons and potentiates spontaneous release in Syt1^+/+^ neurons, without affecting mEPSC amplitudes in all conditions. * p < 0.05, *** p < 0.001, NS p > 0.17 Mann–Whitney U test (b) and ANOVA on ranks (c). The detailed statistical analysis including the numbers of independent experiments is shown in Supplementary Table 1.

## Discussion

In this report we applied the direct classical approach based on testing the effects of a targeted mutation that specifically disrupts a given molecular property - Syt1 oligomerisation - to establish its physiological role in regulation of synaptic neurotransmitter release. We provide several lines of evidence that C2B domain-dependent oligomerisation is essential for Syt1-mediated clamping of both evoked and spontaneous release.

How do Syt1 oligomers could clamp different modes of transmitter release? Besides oligomerisation, binding to SNARE complexes is also necessary for the Syt1 clamping function^12, 26^. A recent X-ray structure has provided an insight into the architecture of pre-fusion Syt1-complexin-SNARE complex^26^. This structure can be combined with the molecular architecture of Syt1 ring-like oligomers into a simple model which could explain the Syt1 clamping function in molecular terms^14^. This model assumes that Syt1 oligomers are formed at the interface between RRP vesicles and the presynaptic membrane and interact with the SNAREpins as visualised in the crystal structure (Supplementary Fig. 4). This Syt1-SNARE organisation would sterically interfere with the completion of SNAREpin zippering thus blocking fusion. The reversal of this fusion clamp – that is, activation of release – would require complete or partial disassembly of Syt1 oligomers which is gated by Ca^2+^ binding by Syt1 C2 domains. This model could provide a plausible explanation for how Ca^2+^-sensitive Syt1 oligomers dynamically regulate different modes of transmitter release triggered not only by Syt1 but also by the other Ca^2+^ release sensors *(e.g.* Syt7, Fig. 2d and Fig. 3c)^1–3, 8, 27^. Apart from the steric occlusion model, Syt1 oligomers are also expected to negatively regulate C2B domain membrane loop insertion and activation of the associated SNAREpins by stabilising the Ca^2+^-binding aliphatic loops at the dimer interface (Fig. 1a). This is also likely to contribute to the gating of different modes of vesicular release. It is worth noting that a substantial proportion spontaneous release is mediated by synaptic vesicles that are molecularly distinct from those that mediate the evoked release^28^. It remains to be determined whether Syt1 contributes to the release of this vesicular pool.

The local concentration of Syt1 on a docked synaptic vesicle is expected to be in the millimolar range (considering 15 – 20 copies of Syt1 localised within 5 nm of synaptic vesicle membrane). The Kd for Syt1 oligomerisation is ∼ 10 μM^29^, so it is reasonable to expect that higher-order Syt1 oligomers will form under these conditions. Furthermore, given the exceedingly high local concentration it is possible that Syt1^F349A^ oligomers may still assemble under synaptic vesicles, though they are expected to have higher dissociation rates and thus be de-stabilised compared to Syt1^WT^. In fact, Syt1^F349A^ does not completely abolish oligomerisation of Syt1^WT^ when mixed together *in vitro* (Fig. 1b). On the other hand, we observe a similar loss-of-clamp phenotype in both rescue and dominant interference experiments in neurons. This argues that formation of oligomers above a certain critical size is likely required for the Syt1-medated clamping function.

Whilst the precise spatial organisation of Syt1 oligomers *in vivo* remains unclear, recent cryo-electron tomography experiments in PC12 cells suggest that fully assembled Syt1 ring-like oligomers may indeed exist on RRP vesicles^30^. These experiments revealed a circular arrangement of six modules of the release machinery under docked synaptic-like vesicles, which is disrupted by overexpression of Syt1^F349A^. This suggests that in addition to the key role for Syt1 oligomers in clamping of different modes of transmitter release (described in the present study), Syt1 oligomerisation could also provide a framework to template and connect multiple SNAREpins to allow their cooperative function.

## Methods

### Lipid Monolayer Assay

Lipid monolayers were set up as described previously^15–17^. Briefly, degassed ultrapure filtered water was injected through a side port to fill up 4-mm diameter and 5-mm depth wells in a Teflon block. The flat-water surface was coated with 0.6 μl of phospholipid mixture PC/PS with molar ratio of 60:40, 0.5 mM in Chloroform. Teflon block was then stored in a sealed humidity chamber for 1 h at room temperature to allow the chloroform to evaporate. Lipid monolayers formed at the air/water interface were then recovered by placing the carbon-coated EM grids (400 mesh; Ted Pella) on top of each water droplet for 1 min. Grids were raised above the surface of the Teflon block by injecting ultrapure H_2_O into the side port and then immediately lifted off the droplet. Protein mixes were diluted to 5 μM in 20 mM 3-(N-morpholino)propanesulfonic acid (pH 7.5), 15 mM KCl, 1 mM EDTA, 2 mM MgAC_2_, 1 mM DTT, and 4% (wt/vol) trehalose buffer, added to the lipid monolayer on the grid and then incubated in a 37 °C humidity chamber for 1 min. The final KCl concentration in the buffer was adjusted to 10 mM. Finally, the dried grids were negatively stained with uranyl acetate solution (1% wt/vol) and examined on an FEI Tecnai T12 microscope operated at 120 kV. The defocus range used for our data was 0.6–2.0 μm. Images were recorded under low-dose conditions (∼20 e^−^/Å2) on a 4K × 4K CCD camera (UltraScan 4000; Gatan, Inc.) at a nominal magnification of 42,000×. Micrographs were binned by a factor of 2 at a final sampling of 5.6 Å per pixel on the object scale. The image analysis, including size distribution measurements, was carried out using ImageJ software (NIH, Bethesda). The following DNA constructs, which have been previously validated in ref.^17^, were used: pCDFDuet-GST-PreScission-Syt1-C2AB-WT and Syt1-C2AB-F349A. The lipids 1-palmitoyl-2-oleoyl-sn-glycero-3-phosphocholine and 1,2-dioleoyl-sn-glycero-3-phospho-L-serine were purchased from Avanti Polar Lipids.

### Neuronal cultures

Experiments conformed to the Animals (Scientific Procedures) Act 1986, and were approved by the ethics committee of the UCL Institute of Neurology. Primary cortical neurons were produced from either Syt1^+/+^ or Syt1^−/−^ mice (B6; 129S-Syt1tm1Sud/J, Jackson Laboratory) and cultured in Neurobasal A/B27-based medium (Thermo Fisher Scientific). Briefly, cortices were dissected from postnatal day 0 (P0) pups and dissociated by enzymatic digestion in 0.25% trypsin for 10 min at 37°C and then triturated using a standard p1000 micropipette. Neurons were plated on poly-L-lysine (1 mg/ml; Sigma-Aldrich) treated 19-mm glass coverslips at a density of 100,000–120,000 cells per coverslip (for imaging and electrophysiological experiments) or on poly-L lysine (0.1 mg/ml)-treated 35-mm plastic dishes at a density of 500,000 cells per dish for Western Blot analysis.

### Construction of plasmids

Syt1-Myc-tagged lentiviral constructs were generated using the pCMV-Myc-N vector (Clontech) containing full-length rat Syt1 and a preprotachykinin signal sequence cloned upstream of the Myc tag as described in ref.^17^ The F349A mutation was introduced using site directed mutagenesis (QuickChange, Agilent Technologies). Myc-tagged Syt1^WT^ and Syt1^F349^ sequences were then subcloned into the lentiviral vector L309 (a gift from T. Südhof, Stanford University^31^) under the ubiquitin promoter using BamHI and BsrGI restriction sites, thereby removing the IRES-EGFP region. For Syt7 KD experiments, the previously validated oligonucleotide sequence KD606 (AAAGACAAGCGGGTAGAGAAA)^32^ was cloned under the U6 promoter present in L309-Myc-Syt1 constructs using AscI and PacI restriction sites. To generate the mCherry-tagged Syt1 constructs, the pcDNA3 pHluorin rat Syt1 plasmid (a gift from V. Haucke^33^, Leibniz-Forschungsinstitut für Molekulare Pharmakologie, Berlin) was modified by exchanging the pHluorin and mCherry sequences using HindIII and AgeI restriction sites. mCherry-tagged Syt1 WT and F349A sequences were then moved into the pCCL lentiviral backbone (a gift from S. Schorge, University College London) under the human Synapsin promoter using BamHI and SalI restriction sites. The pFU_SypHy plasmid was kindly provided by A. Maximov (The Scripps Research Institute, La Jolla). The pAAV.hSynap.SF-iGluSnFR.A184V plasmid^25^ was kindly provided by J. Marvin and L. Looger (Janelia Research Campus).

### Lentiviral production and transduction of primary cortical cultures

The production of vesicular stomatitis virus-pseudotyped second and third-generation lentiviruses was performed by co-transfection of the pFUGW/L309/PCCL based expression vectors and two (pCMVdelta and pCMV-VSV-G) or three helper plasmids (pCMV-VSV-G, pMDLg/pRRE, pRSV-Rev) in human embryonic kidney 293T (HEK293T) cells using Lipofectamine 2000 (Thermo Fisher Scientific) following the instructions of the manufacturer. Primary neurons were transduced at 7 days *in vitro* (DIV) using 10 multiplicity of infection either with viruses expressing the Myc-tagged Syt1^WT^ and Syt1^F349A^ (Fig.1 and Fig.3) or the mCherry-tagged Syt1^WT^ and Syt1^F349A^. Transduction efficiency (80-95% range) was verified using immunofluorescence analysis (see also Extended Data Figs. 2 – 5). For experiments in Fig.1 neuronal cultures were also transduced with the sypHy expressing lentivirus. For experiments in Fig.2 cortical neurons were transfected at 5 DIV with pAAV.hSynap.SF-iGluSnFR.A184V plasmid using Neuromag reagent (#KC30800, OZ Biosciences). This allowed expression of the iGluSnFR probe only in a small (∼ 3 – 5 %) subpopulation of neurons which was essential for imaging of vesicular release in individual synaptic boutons.

### SypHy imaging experiments

All imaging and electrophysiological experiments were conducted in a custom-made open laminar flow field–stimulation chamber (0.35 ml) at 23–25 °C and 15–22 DIV. The Extracellular Buffer (EB1) contained in (mM): NaCl (125), KCl (2.5), MgCl_2_ (2), CaCl_2_ (2), glucose (30), HEPES (25), pH 7.4^34, 35^. To block recurrent activity EB1 was supplemented with NBQX (10 μM, Ascent Scientific) and DL-AP5 (50 μM, Ascent Scientific). SypHy imaging was performed on an inverted Zeiss Axiovert 200 fluorescence microscope using a 63× (1.4 NA) oil-immersion objective. Images (12 bit) were acquired using a Prime95B CMOS camera (Photometrics) coupled to a 488 nm excitation light emitting diode (LED) light source and a 510 long-pass emission filter, with an exposure time of 50ms. Evoked sypHy responses were recorded in an ∼ 180 × 180-μm region of interest (ROI) typically containing 50 – 200 individual boutons. Images were analysed using ImageJ (NIH, Bethesda) and MATLAB (MathWorks) custom-made software scripts. Synaptic boutons were identified by subtracting the resting sypHy fluorescence from the peak fluorescence after 20 action potentials 100 Hz burst, which was previously shown to trigger release of the entire RRP of vesicles^20^. This was followed by the generation of a binary mask that was used to measure the average fluorescence response in all identified boutons in each experiment (Fig.1c). After subtracting the background, the data were normalised to the resting sypHy signal. The release probability of RRP vesicles was estimated by normalising the average (10 sweeps) sypHy response to a single spike to the response to 20 action potentials 100 Hz train^20^: pv = ΔF_1AP_/ΔF_20AP_. We also used the increase of sypHy fluorescence measured 2 sec after the action potential burst as a lower estimate of the asynchronous release (ΔF_Asynch_). Potentially ΔF_Asynch_ could be affected by synaptic vesicle endocytosis and re-acidification. However, based on the measurements of vesicular endocytosis and re-acidification rates in similar experimental conditions we estimate that the possible error should not exceed ∼ 10–15 %^36, 37^.

### IGluSnFR imaging experiments

iGluSnFR imaging was performed on an inverted Olympus IX71 fluorescence microscope equipped with a 60× (1.35 NA) oil-immersion objective and a QuantEM 512SC EM-CCD camera (Photometrics). iGluSnFR fluorescence was recoded using a 470 nm excitation LED and a 500 – 550 nm band-pass emission filter. Expression of the mCherry tagged Syt1^WT^ and Syt1^F349A^ constructs was confirmed using a white LED combined with a 535 – 560 nm band-pass excitation filter and a 570 nm long-pass emission filter. The carbonated Extracellular Buffer 2 (EB2) contained in (mM): NaCl (125), NaHCO_3_ (26), NaHPO_4_ (1.25), KCl (2.5), CaCl_2_ (2), MgCl_2_ (1.3) and Glucose (12); NBQX (10 μM), DL-AP5 (50 μM) and picrotoxin (50 μM, Tocris Bioscience) were added to block the recurrent activity.

An isolated putative pyramidal-like neuron expressing iGluSnFR was whole-cell patch-clamped with a 5-7 MΩ fire-polished borosilicate glass pipette (current-clamp configuration) using the following intracellular solution (in mM): K+-gluconate (105), KCl (30), HEPES (10), phosphocreatine Na_2_ (10), ATP-Mg (4), GTP-NaH_2_O (0.3) and 1 EGTA, balanced to pH 7.3 with KOH, −10 mV liquid junction potential. Neurons were stimulated using 15 ms current pulses (∼300 pA), generating a total of 51 action potentials at 5Hz. Action potential waveform recordings were obtained with an Axon Multiclamp 700B amplifier, CV-7 headstage (Molecular Devices). Signal was acquired at 20 KHz (4 KHz Bessel-filtering), with WinWCP software controlling a National Instrument board (NI USB-6221). Candidate presynaptic boutons were selected for imaging by tracing the axon from the patched neuron and identifying regions with stereotypical ∼ 1-2 μm bulbous structures, usually within 200 μM distance from the soma. A ROI of ∼ 137×20 μm (512×75 pixels) was imaged with 6.05 ms temporal resolution, using 150 EM gain. Image acquisition was performed using Micro Manager software^38^. For each cell, from one to five different ROIs were recorded. Imaging and electrophysiological recordings were synchronised, allowing us to identify the precise timing of release event with respect to the presiding action potential. The acquired image stack was first bandpass filtered at 0.5-30Hz (z axis), then median filtered (x and y axis, 3×3 pixels), revealing the location of the stereotypical iGluSnFR responses (see Supplementary Movie 1). Next, a maximal projection was produced from the filtered image stack, from which ROIs corresponding to putative boutons were selected using ImageJ software. This set of ROIs were used to extract iGluSnFR responses from the raw, unfiltered image.

To determine the timings and the amplitudes of quantal releases in each bouton the average unitary iGluSnFR response (defined as an instantaneous rise and an exponential decay f_unitary_(t)=exp^−t/T^, where t = 60 ms is the experimentally determined decay time constant) was deconvolved from the raw iGluSnFR response as: F^−1^[F(f_iGluSnFR_(t))/F(f_unitaty_(t))], where F is the discrete Fourier transform and F^−1^ is the inverse Fourier transform (MATLAB). Events were defined as synchronous if they occurred within 3 frames (∼18 ms) of the preceding action potential, and asynchronous in any other condition. Amplitude was estimated in quanta, which was calculated as: (i) the average 5 smallest synchronous events for high release probability boutons; (ii) the average 2 smallest synchronous events for low release probability boutons or (iii) the smallest event if no 2 synchronous releases were detected.

### mEPSC recording and analysis

Spontaneous mEPSCs were recorded in EB1 supplemented with 1μM tetrodotoxin (Tocris Bioscience), 100μM picrotoxin and 1μM CGP55845 (Tocris Bioscience). Patch pipettes (8-12 MΩ) were fabricated from borosilicate glass capillaries using a micropipette puller (P-97, Sutter). The intracellular solution contained (in mM) CS-OH (125), Gluconic acid (125), HEPES (10), Na-Phosphocreatine (10), NaCl (8), Na-GTP (0.33), Mg-ATP (4), EGTA (0.2), TEA-Cl (5), pH 7.33. Putative pyramidal cells were visualised using differential interference contrast and an inverted microscope (Axio Observer A1, Zeiss) equipped with a CMOS Prime95B camera. Postsynaptic mEPSCs were recorded in voltage clamp mode at a holding potential of −65mV using a MultiClamp 700B (Axon Instruments), filtered at 3kHz (Bessel filter) and digitised at 10kHz using NIDAQ-MX National Instruments Interface Card and the WinEDR software (Strathclyde Electrophysiology). No correction was made for the junction potential between the patch pipette intracellular and extracellular solutions. Series resistance (Rs) was monitored every ∼ 2 mins with 10 mV hyperpolarizing steps and was not corrected. No compensation was made for whole cell (Cm) and electrode capacitance (Cp). To confirm the AMPAR origin of the detected events in a subset of cells the AMPA/Kainate receptor antagonist NBQX (25 μM) was bath applied at the end of experiments. Individual mEPSC detection was performed in a single time window (30-120s) at least two minutes after obtaining whole cell configuration, to allow for stable recording conditions. Recordings were band-pass filtered between 4 Hz and 5KHz (pClamp 10, Molecular Devices). Threshold criteria for mEPSC detection were: amplitude > 5 pA, duration 0.5-100 ms, with noise rejection 0.5 ms. Recordings were excluded if holding current exceeded −350 pA, Rs exceeded 35 MΩ, and/or Rs varied by >20 % during the recording.

### Immunofluorescence analysis

Neurons were fixed with 4% paraformaldehyde in PBS for 10 minutes and permeabilised with 0.1% Triton X-100 in PBS (Phosphate-Buffered Saline, Thermo Fisher Scientific) for 10 minutes. Nonspecific binding sites were blocked by incubation for 30 minutes with 3% BSA (bovine serum albumin) and 0.1% Triton X-100 in PBS before 2 hrs incubation at room temperature with primary antibodies. Neurons were washed 3 times with PBS and incubated with secondary antibodies conjugated with Alexa Fluor (488, 555 or 568) for 45 minutes at RT. Coverslips were washed 3 times in PBS and mounted using a mounting media containing DAPI (Thermo Fisher, P36931) to visualise nuclei. For Syt7 KD experiments, cortical neurons were fixed with cold methanol for 7 minutes and then processed as described above without any permeabilization step. The following primary antibodies were used: anti-Syt1 (105 003, Synaptic Systems); anti-Myc (MA1-980, Thermo Fisher), anti-Map2 (188-004, Synaptic Systems); anti-Syt7 (105 173, Synaptic Systems); anti-VAMP2 (104-202, Synaptic Systems). Somatic and synaptic fluorescence intensity were quantified using ImageJ (NIH, Bethesda). Somatic fluorescence intensity was estimated by manual tracking of cell bodies while synaptic intensity was quantified by applying 2μm diameter circular ROIs over the identified synaptic boutons. Synaptic density was estimated by manual count of VAMP2-positive boutons along selected Map2 dendritic branches. For Syt7 KD experiments, a binary mask created from the Myc staining image (green channel) was applied to estimate the total Syt7 intensity (red channel).

### Western blotting

Cortical neurons at 16 DIV were harvested in RIPA lysis buffer (50 mM Tris-HCl pH 7.5, 150 mM NaCl, 1% NP-40, 0.5% sodium deoxycholate, 0.1% SDS, 1 mM EDTA, 1 mM EGTA) containing Halt™ protease inhibitor cocktail (1 in 100; ThermoFisher). After centrifugation at 16,000 × g for 15 min at 4°C, samples were loaded on SDS-PAGE gel, transferred to nitrocellulose membranes, and immunoblotted with primary antibodies followed by HRP-conjugated secondary antibodies. The following primary antibodies were used: anti-Syt1 (105 003, Synaptic Systems), anti-Syt7 (105 173, Synaptic Systems) and anti-GAPDH (G8795, Sigma). Immunoreactive bands were visualized by enhanced chemiluminescence (ECL), captured using a Biorad ChemiDoc reader and analysed using ImageLab software. Syt1 and Syt7 protein levels were normalized to the GAPDH loading control.

### Statistical analysis

The distribution of data in each set of experiments was first tested for normality using the Shapiro-Wilk test. The similarity of variances between each group of data was tested using the F test. Normally distributed data were presented as mean ± s.e.m., each plot also contained the individual data points. Student’s t tests for group means or one way ANOVA were used as indicated. The data sets that failed the normality test were presented using box-and-whisker plots (box 25^th^ – 75^th^ percentiles, whiskers 10^th^ – 90^th^ percentiles), each plot also contained the individual data points. Here Mann-Whitney U test and ANOVA on ranks were used as indicated. The detailed statistical analysis is presented in Extended Data Table 1 and Extended Data Table 2. No statistical methods were used to pre-determine sample sizes, but our sample sizes are similar to those reported in previous publications in the field^12, 20, 26, 35^. In Figure 1 data analysis was performed blind to the conditions. In Figures 2 and 3 both data collection and analysis were performed blind to the conditions. All statistical tests were performed using SigmaPlot 11 (Systat Software).

## Author contributions

E.T., J.E.R., S.S.K. and K.E.V. designed the study. E.T. performed sypHy imaging experiments, preparation / viral transduction of neuronal cultures and immunocytochemistry. O.D.B. performed *in vitro* analysis of Syt1 oligomerisation. P.R.F.M. performed iGluSnFR imaging experiments. D.K. and E.N. performed electrophysiological recordings. J.C. assisted with construction of recombinant plasmids. Y.T. and K.E.V contributed to the analysis of sypHy and iGluSnFR imaging experiments. S.S.K and K.E.V wrote the manuscript.

## Acknowledgements

We are grateful to D. M. Kullmann for reading the manuscript and providing critical feedback. The study was supported by the Wellcome Trust (K.E.V. and J.E.R.), Medical Research Council (K.E.V.) and the NIH grant DK027044 (J.E.R.).

