## Supplemental Figures for "Synaptotagmin 1 oligomers clamp and regulate different modes of neurotransmitter release"

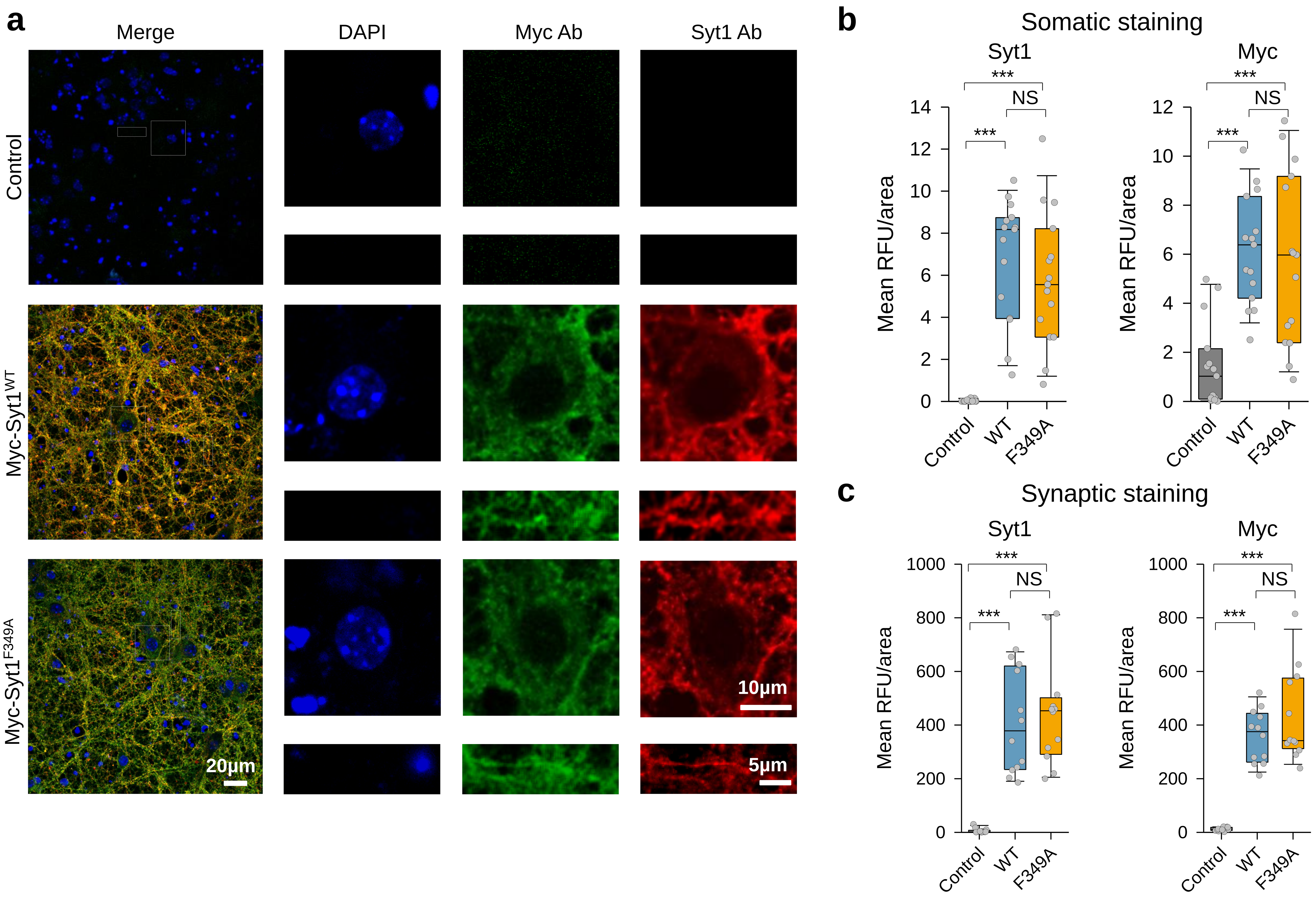


**Supplementary Figure 1.**

**Overexpression of Myc-Syt1$^{WT}$ and Myc-Syt1$^{F349A}$ in Syt1$^{-/-}$ neurons.**

(**a**) Representative images of untransduced Syt1$^{-/-}$ neurons (Control) and of Syt1$^{-/-}$ neurons transduced either with Myc-Syt1$^{WT}$ or Myc-Syt1$^{F349A}$ lentiviruses immunostained with Abs against Myc tag (green) and Syt1 (red). DAPI was used to visualise the cell nuclei. Two different types of ROIs (right panels and corresponding white boxes on the original images) containing either somas or neurites were used to quantify somatic and synaptic Myc and Syt1 immunoreactivity respectively. Transduction efficiency was in the range of 80 – 95%.

(**b**, **c**) Quantification of somatic and synaptic Myc and Syt1 immunofluorescence levels showing rescue of Syt1 expression in the transduced neurons. *** $p < 0.001$, NS $p > 0.3$, Mann–Whitney U test. The detailed statistical analysis is reported in Supplementary Table 1.

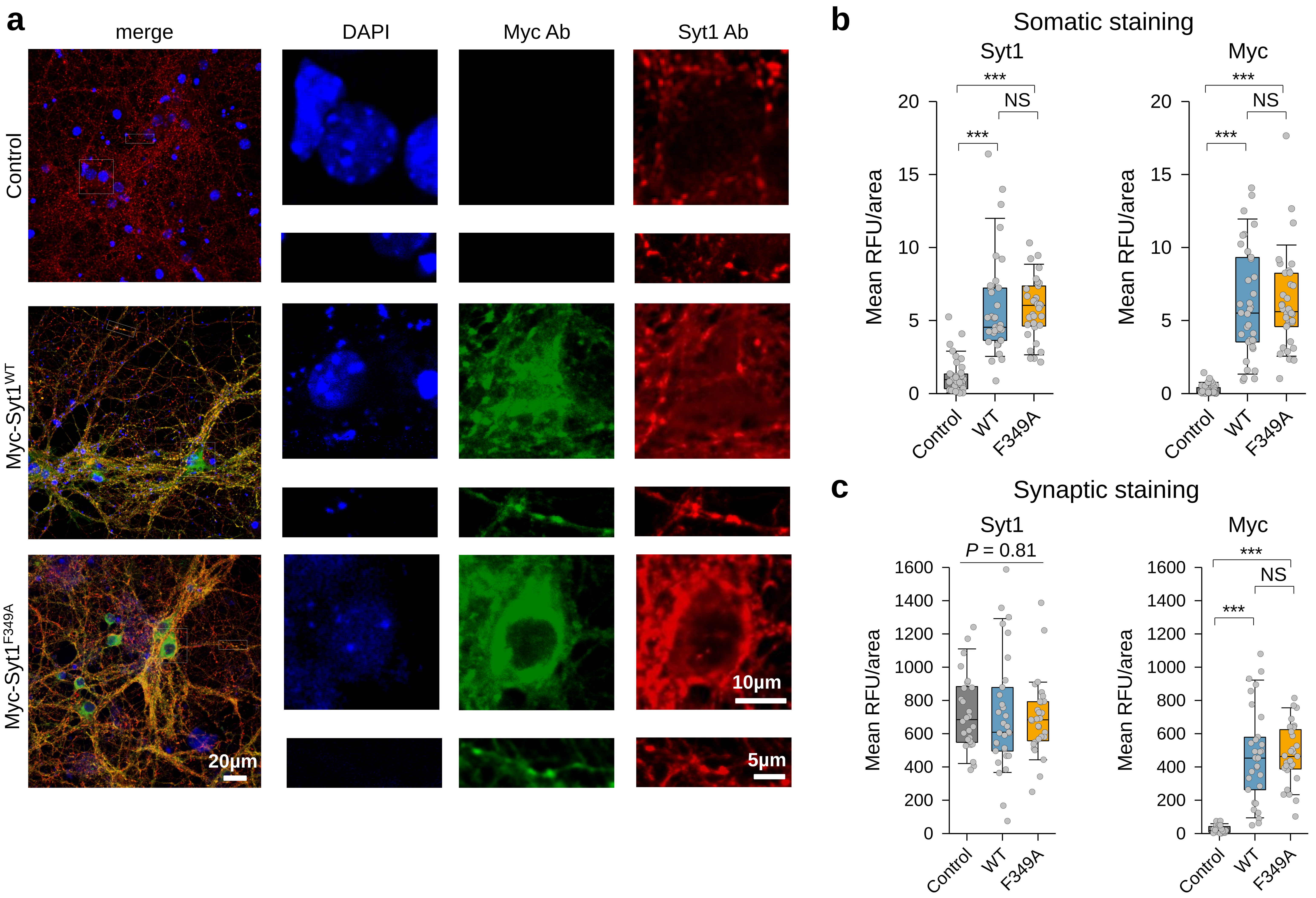


**Supplementary Figure 2.**

**Overexpression of Myc-Syt1$^{WT}$ and Myc-Syt1$^{F349A}$ in Syt1$^{+/+}$ neurons.**

(**a**) Representative images of untransduced Syt1$^{+/+}$ neurons (Control) and of Syt1$^{+/+}$ neurons transduced either with Myc-Syt1$^{WT}$ or Myc-Syt1$^{F349A}$ lentiviruses immunostained with Abs against Myc tag (green) and Syt1 (red). DAPI was used to visualise the cell nuclei. Two different types of ROIs (right panels and corresponding white boxes on the original images) containing either somas or neurites were used to quantify somatic and synaptic Myc and Syt1 immunoreactivity respectively. Transduction efficiency was > 90 – 95 %

(**b**, **c**) Quantification of somatic and synaptic Myc and Syt1 immunofluorescence levels showing increase of Syt1 expression in somatic but not synaptic areas of the transduced neurons. *** $p < 0.001$, NS $p > 0.5$, Mann–Whitney U test. The detailed statistical analysis is reported in Supplementary Table 1.

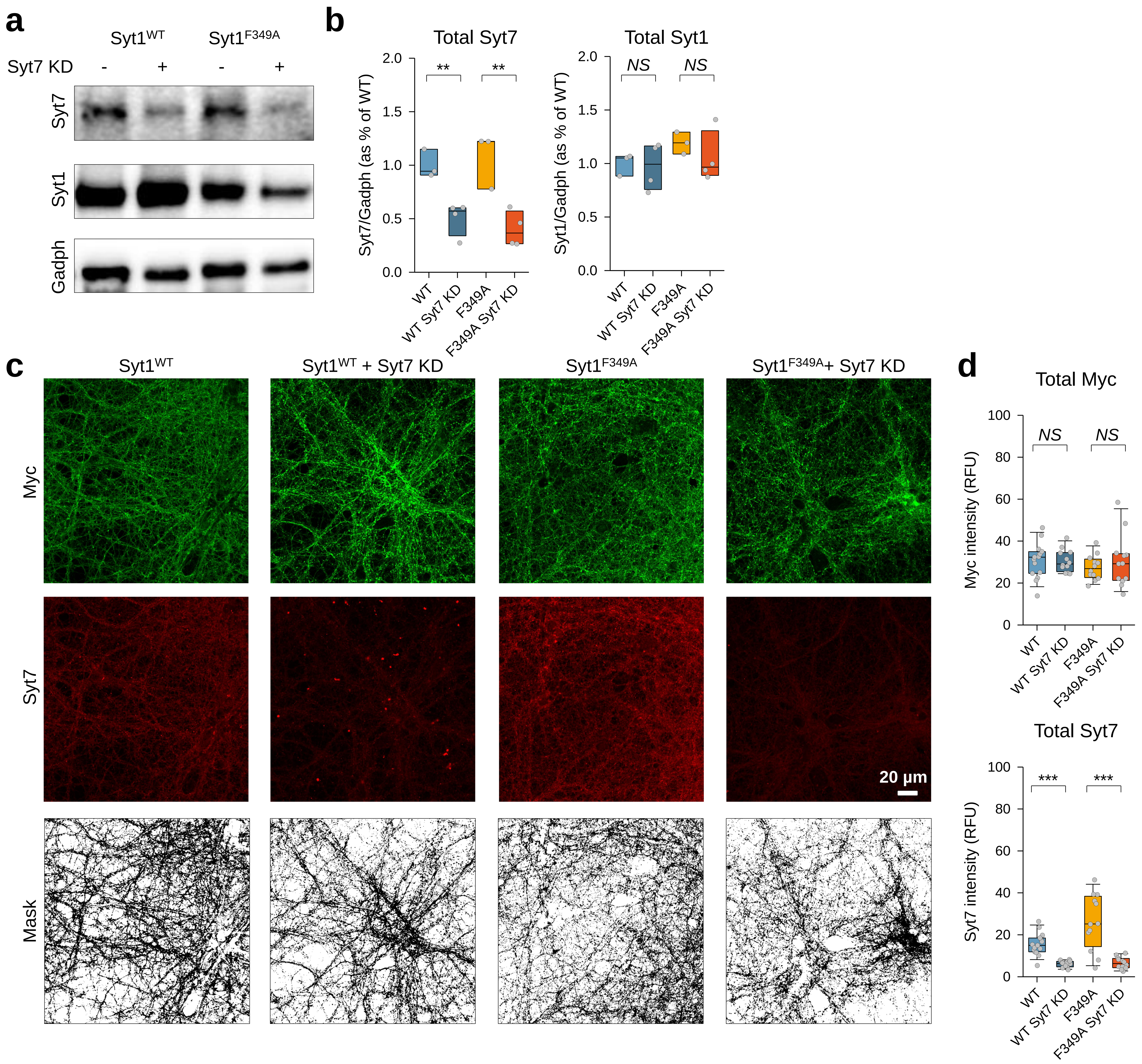


**Supplementary Figure 3.**

**Syt7 KDs in Myc-Syt1$^{WT}$ and Myc-Syt1$^{F349A}$ expressing Syt1$^{+/+}$ neurons.**

(**a**) Representative Western Blot analysis of Syt7 and Syt1 protein levels in cell lysates from Syt1$^{+/+}$ neurons transduced with Myc-Syt1$^{WT}$, Myc-Syt1$^{F349A}$, Myc-Syt$^{WT}$-Syt7-KD or Myc-Syt1$^{F349A}$-Syt7-KD lentiviral constructs.

(**b**) Quantification of Syt7 and Syt1 protein levels normalised to the Gadph loading control.

(**c**) Representative images of Syt1$^{+/+}$ neurons transduced with Myc-Syt1$^{WT}$, Myc-Syt1$^{F349A}$, Myc-Syt$^{WT}$-Syt7-KD or Myc-Syt1$^{F349A}$-Syt7-KD lentiviral constructs, immunostained with Abs against Myc tag (green) and Syt7 (red). A binary mask created from the Myc image was applied to estimate the relative Syt7 immunoreactivity.

(**d**) Quantification of total Myc and Syt7 fluorescence reveals a significant decrease of Syt7 levels in Syt7 KD neurons compared to the relative controls. ** $p < 0.01$, *** $p < 0.001$, NS $p > 0.4$, Student t-test (**b**) and Mann–Whitney U test (**d**). The detailed statistical analysis is reported in Supplementary Table 1.

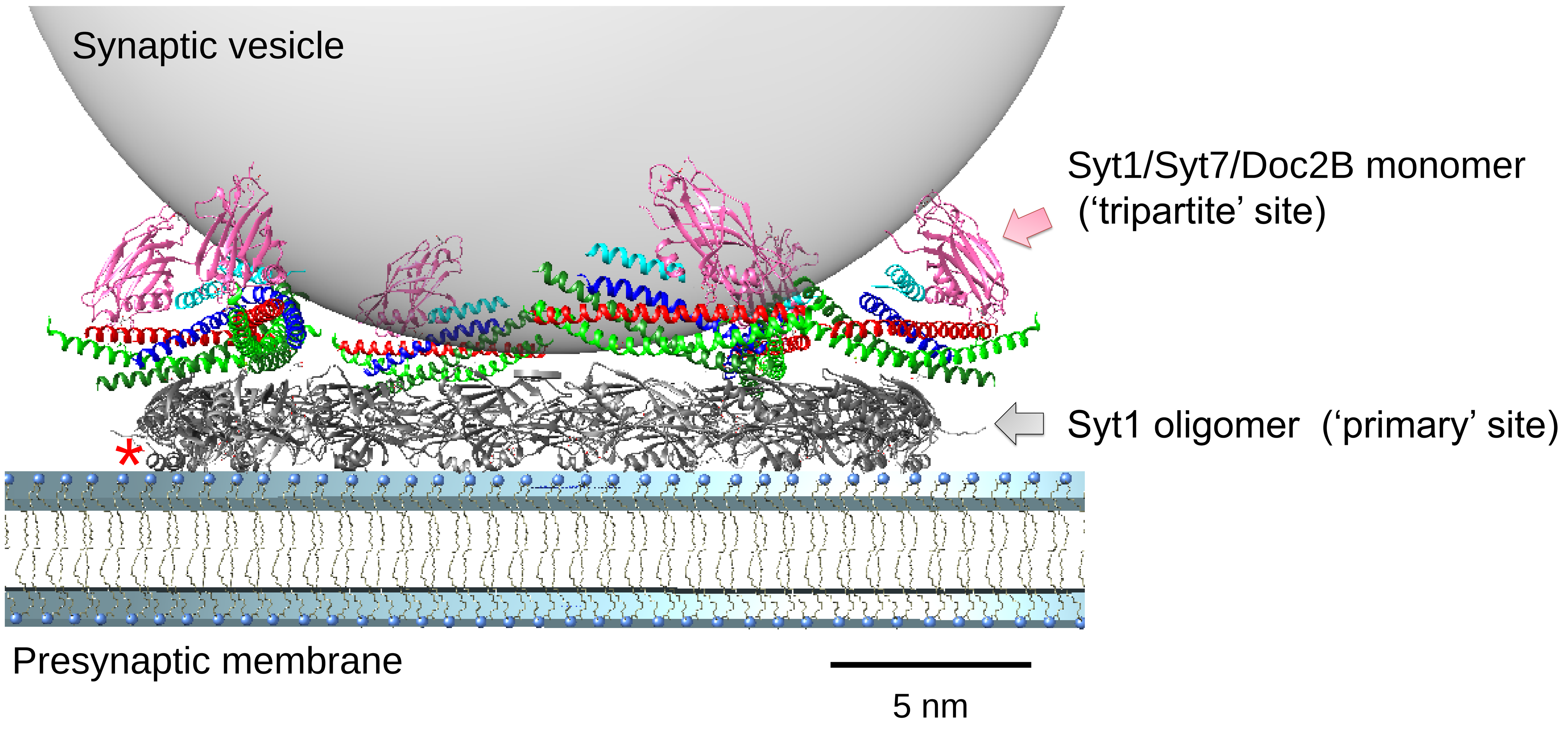


**Supplementary Figure 4.**

**Molecular model for the clamping of vesicular fusion by Syt1 ring-like oligomers**

This model, combining the recent X-ray SNAREpin-Syt1-complexin structure[26] with the molecular architecture of Syt1 ring-like oligomers, could potentially explain the Syt1 clamping function in molecular terms[14]. The X-ray crystal structure shows that each pre-fusion partially assembled SNARE complex (SNAREpin) contains two distinct Syt1 binding sites: a complexin-independent 'primary' site, which can only be occupied by synaptotagmins that mediate synchronous release (Syt1, Syt2 and Syt9) and a complexin-dependent 'tripartite' site which is potentially available to all synaptotagmin isoforms, including Syt7[26]. We posit that the full complement (15-20 copies) of Syt1 on the vesicle oligomerise at the site of docking triggered by PIP2 binding. This Syt1 oligomer (grey) is then ideally positioned to simultaneously template multiple SNAREpins via the 'primary' binding motif, which is accessible and free to interact in the ring oligomer. Note that in the ring-oligomer configuration, the conserved helical extension (red asterisk) involved in the tripartite binding locates towards the membrane and is thus unavailable. The height of Syt1 oligomers combined with the SNAREs atop would maintain the two bilayers far apart (~4 nm) to allow the N-terminal assembly but to sterically block the complete assembly of SNAREs. Furthermore, Syt1 oligomers are also expected to restrain the bound SNAREpins from moving inward toward the incipient fusion pore. Thus, the ring oligomer will clamp the synaptic vesicle fusion. In this arrangement, a second independent C2B domain (magenta) from either Syt1, Syt7 or Doc2B[26] can bind the SNAREpin via the 'tripartite' site in conjunction with complexin (cyan) and further, stabilise the fusion clamp. Upon $Ca^{2+}$ influx, the Syt1 ring oligomers are disrupted as Syt1 molecules rotate to insert into the plasma membrane. This frees the SNAREpins to complete zippering and trigger fusion. In this manner, the Syt1 oligomer could mediate a $Ca^{2+}$-sensitive clamp on vesicular fusion. Critically, this dual-clamp model illustrates how Syt1 oligomerization could gate activation of both Syt1 and the second $Ca^{2+}$ sensor (e.g. Syt7). Adapted from ref.[14].
