## Supplemental Table 1 for "Synaptotagmin 1 oligomers clamp and regulate different modes of neurotransmitter release"

### Supplementary Table 1 Statistical Analysis

#### Figure 1

| *Condition* | *Number of preparations* |
|---|---|
| $Syt1^{WT}$ | *3* |
| $Syt1^{F349A}$ | *3* |
| $Syt1^{WT}$ + $Syt1^{F349A}$ | *3* |

**Figure 1d Total oligomeric structures**

| *Condition 1* | *Condition 2* | *P value* | *Statistical test* |
|---|---|---|---|
| $Syt1^{WT}$ | $Syt1^{F349A}$ | P < 0.001 | Student t-test |
| $Syt1^{WT}$ | $Syt1^{WT}$ + $Syt1^{F349A}$ | P = 0.020 | Student t-test |
| $Syt1^{F349A}$ | $Syt1^{WT}$ + $Syt1^{F349A}$ | P < 0.001 | Student t-test |

**Figure 1d Ring-like oligomeric structures**

| *Condition 1* | *Condition 2* | *P value* | *Statistical test* |
|---|---|---|---|
| $Syt1^{WT}$ | $Syt1^{F349A}$ | P < 0.001 | Student t-test |
| $Syt1^{WT}$ | $Syt1^{WT}$ + $Syt1^{F349A}$ | P < 0.001 | Student t-test |
| $Syt1^{F349A}$ | $Syt1^{WT}$ + $Syt1^{F349A}$ | P = 0.048 | Student t-test |

**Figure 1e Oligomeric rings diameter**

| *Condition 1* | *Condition 2* | *P value* | *Statistical test* |
|---|---|---|---|
| $Syt1^{WT}$ | $Syt1^{F349A}$ | P = 0.196 | Student t-test |
| $Syt1^{WT}$ | $Syt1^{WT}$ + $Syt1^{F349A}$ | P < 0.001 | Student t-test |
| $Syt1^{F349A}$ | $Syt1^{WT}$ + $Syt1^{F349A}$ | P < 0.001 | Student t-test |

#### Figure 2

| *Condition* | *Number of cells* | *Number of preparations* |
|---|---|---|
| $Syt1^{-/-}$ Control | *12* | *6* |
| $Syt1^{-/-}$ $Syt1^{WT}$ | *15* | *7* |
| $Syt1^{-/-}$ $Syt1^{F349A}$ | *15* | *7* |
| $Syt1^{+/+}$ Control | *11* | *3* |
| $Syt1^{+/+}$ $Syt1^{WT}$ | *23* | *9* |
| $Syt1^{+/+}$ $Syt1^{F349A}$ | *25* | *10* |
| $Syt1^{+/+}$ $Syt1^{WT}$ Syt7 KD | *24* | *10* |
| $Syt1^{+/+}$ $Syt1^{F349A}$ Syt7 KD | *28* | *11* |

**Figure 2b**

Syt1$^{-/-}$ background, one way ANOVA on ranks P = 0.981

Syt1$^{+/+}$ background, one way ANOVA on ranks P = 0.224

**Figure 2c**

| *Condition 1* | *Condition 2* | *P value* | *Statistical test* |
|---|---|---|---|
| Syt1$^{-/-}$ Control | Syt1$^{-/-}$ Syt1$^{WT}$ | P = <0.001 | Mann-Whitney U test |
| Syt1$^{-/-}$ Control | Syt1$^{-/-}$ Syt1$^{F349A}$ | P = <0.001 | Mann-Whitney U test |
| Syt1$^{-/-}$ Syt1$^{WT}$ | Syt1$^{-/-}$ Syt1$^{F349A}$ | P = <0.001 | Mann-Whitney U test |
| Syt1$^{+/+}$ Control | Syt1$^{+/+}$ Syt1$^{WT}$ | P = 0.912 | Mann-Whitney U test |
| Syt1$^{+/+}$ Control | Syt1$^{+/+}$ Syt1$^{F349A}$ | P = 0.007 | Mann-Whitney U test |
| Syt1$^{+/+}$ Syt1$^{WT}$ | Syt1$^{+/+}$ Syt1$^{F349A}$ | P = 0.001 | Mann-Whitney U test |
| Syt1$^{+/+}$ Syt1$^{WT}$ Syt7 KD | Syt1$^{+/+}$ Syt1$^{F349A}$ Syt7 KD | P = 0.007 | Mann-Whitney U test |

**Figure 2d**

| *Condition 1* | *Condition 2* | *P value* | *Statistical test* |
|---|---|---|---|
| Syt1$^{-/-}$ Control | Syt1$^{-/-}$ Syt1$^{WT}$ | P = 0.020 | Mann-Whitney U test |
| Syt1$^{-/-}$ Control | Syt1$^{-/-}$ Syt1$^{F349A}$ | P = 0.608 | Mann-Whitney U test |
| Syt1$^{-/-}$ Syt1$^{WT}$ | Syt1$^{-/-}$ Syt1$^{F349A}$ | P = 0.020 | Mann-Whitney U test |
| Syt1$^{+/+}$ Control | Syt1$^{+/+}$ Syt1$^{WT}$ | P = 0.854 | Mann-Whitney U test |
| Syt1$^{+/+}$ Control | Syt1$^{+/+}$ Syt1$^{F349A}$ | P = 0.004 | Mann-Whitney U test |
| Syt1$^{+/+}$ Syt1$^{WT}$ | Syt1$^{+/+}$ Syt1$^{F349A}$ | P = 0.003 | Mann-Whitney U test |
| Syt1$^{+/+}$ Syt1$^{WT}$ Syt7 KD | Syt1$^{+/+}$ Syt1$^{F349A}$ Syt7 KD | P = 0.539 | Mann-Whitney U test |

### Figure 3

| *Condition* | *Number of cells* | *Number of preparations* |
|---|---|---|
| $Syt1^{+/+}$ $Syt1^{WT}$ | *17* | *7* |
| $Syt1^{+/+}$ $Syt1^{F349A}$ | *17* | *7* |

**Figure 3c Synchronous release**

| *Condition 1* | *Condition 2* | *P value* | *Statistical test* |
|---|---|---|---|
| $Syt1^{+/+}$ $Syt1^{WT}$ $1^{st}$ AP | $Syt1^{+/+}$ $Syt1^{F349A}$ $1^{st}$ AP | P = 0.084 | Mann-Whitney U test |
| $Syt1^{+/+}$ $Syt1^{WT}$ all APs | $Syt1^{+/+}$ $Syt1^{F349A}$ all APs | P = 0.705 | Mann-Whitney U test |

**Figure 3c Asynchronous release**

| *Condition 1* | *Condition 2* | *P value* | *Statistical test* |
|---|---|---|---|
| $Syt1^{+/+}$ $Syt1^{WT}$ $1^{st}$ AP | $Syt1^{+/+}$ $Syt1^{F349A}$ $1^{st}$ AP | P = 0.988 | Mann-Whitney U test |
| $Syt1^{+/+}$ $Syt1^{WT}$ all APs | $Syt1^{+/+}$ $Syt1^{F349A}$ all APs | P = 0.012 | Mann-Whitney U test |

### Figure 4

| *Condition* | *Number of cells* | *Number of preparations* |
|---|---|---|
| $Syt1^{-/-}$ Control | *25* | *4* |
| $Syt1^{-/-}$ $Syt1^{WT}$ | *20* | *5* |
| $Syt1^{-/-}$ $Syt1^{F349A}$ | *30* | *7* |
| $Syt1^{+/+}$ Control | *21* | *5* |
| $Syt1^{+/+}$ $Syt1^{WT}$ | *21* | *5* |
| $Syt1^{+/+}$ $Syt1^{F349A}$ | *19* | *5* |

**Figure 4b**

| *Condition 1* | *Condition 2* | *P value* | *Statistical test* |
|---|---|---|---|
| $Syt1^{-/-}$ Control | $Syt1^{-/-}$ $Syt1^{WT}$ | P = <0.001 | Mann-Whitney U test |
| $Syt1^{-/-}$ Control | $Syt1^{-/-}$ $Syt1^{F349A}$ | P = 0.240 | Mann-Whitney U test |
| $Syt1^{-/-}$ $Syt1^{WT}$ | $Syt1^{-/-}$ $Syt1^{F349A}$ | P = 0.012 | Mann-Whitney U test |

| | | | |
|---|---|---|---|
| $Syt1^{+/+}$ Control | $Syt1^{+/+}$ $Syt1^{WT}$ | P = 0.814 | Mann-Whitney U test |
| $Syt1^{+/+}$ Control | $Syt1^{+/+}$ $Syt1^{F349A}$ | P = 0.032 | Mann-Whitney U test |
| $Syt1^{+/+}$ $Syt1^{WT}$ | $Syt1^{+/+}$ $Syt1^{F349A}$ | P = 0.021 | Mann-Whitney U test |

**Figure 4c**

$Syt1^{-/-}$ background, one way ANOVA on ranks P = 0.18

$Syt1^{+/+}$ background, one way ANOVA on ranks P = 0.17

### Supplementary Figure 1

| *Condition* | *Number of cells* | *Number of preparations* |
|---|---|---|
| $Syt1^{-/-}$ Control | *12-15* | *3* |
| $Syt1^{-/-}$ $Syt1^{WT}$ | *12-15* | *3* |
| $Syt1^{-/-}$ $Syt1^{F349A}$ | *12-15* | *3* |

**Supplementary Figure 1b anti-Syt1 Ab**

| *Condition 1* | *Condition 2* | *P value* | *Statistical test* |
|---|---|---|---|
| $Syt1^{-/-}$ Control | $Syt1^{-/-}$ $Syt1^{WT}$ | P = <0.001 | Mann-Whitney U test |
| $Syt1^{-/-}$ Control | $Syt1^{-/-}$ $Syt1^{F349A}$ | P = <0.001 | Mann-Whitney U test |
| $Syt1^{-/-}$ $Syt1^{WT}$ | $Syt1^{-/-}$ $Syt1^{F349A}$ | P = 0.300 | Mann-Whitney U test |

**Supplementary Figure 1b anti-Myc Ab**

| *Condition 1* | *Condition 2* | *P value* | *Statistical test* |
|---|---|---|---|
| $Syt1^{-/-}$ Control | $Syt1^{-/-}$ $Syt1^{WT}$ | P = <0.001 | Mann-Whitney U test |
| $Syt1^{-/-}$ Control | $Syt1^{-/-}$ $Syt1^{F349A}$ | P = <0.001 | Mann-Whitney U test |
| $Syt1^{-/-}$ $Syt1^{WT}$ | $Syt1^{-/-}$ $Syt1^{F349A}$ | P = 0.590 | Mann-Whitney U test |

**Supplementary Figure 1c anti-Syt1 Ab**

| *Condition 1* | *Condition 2* | *P value* | *Statistical test* |
|---|---|---|---|
| $Syt1^{-/-}$ Control | $Syt1^{-/-}$ $Syt1^{WT}$ | P = <0.001 | Mann-Whitney U test |

| | | | |
|---|---|---|---|
| Syt1$^{-/-}$ Control | Syt1$^{-/-}$ Syt1$^{F349A}$ | P = <0.001 | Mann-Whitney U test |
| Syt1$^{-/-}$ Syt1$^{WT}$ | Syt1$^{-/-}$ Syt1$^{F349A}$ | P = 0.583 | Mann-Whitney U test |

**Supplementary Figure 1c anti-Myc Ab**

| *Condition 1* | *Condition 2* | *P value* | *Statistical test* |
|---|---|---|---|
| Syt1$^{-/-}$ Control | Syt1$^{-/-}$ Syt1$^{WT}$ | P = <0.001 | Mann-Whitney U test |
| Syt1$^{-/-}$ Control | Syt1$^{-/-}$ Syt1$^{F349A}$ | P = <0.001 | Mann-Whitney U test |
| Syt1$^{-/-}$ Syt1$^{WT}$ | Syt1$^{-/-}$ Syt1$^{F349A}$ | P = 0.371 | Mann-Whitney U test |

### Supplementary Figure 2

| *Condition* | *Number of cells* | *Number of preparations* |
|---|---|---|
| Syt1$^{+/+}$ Control | *26-39* | *6* |
| Syt1$^{+/+}$ Syt1$^{WT}$ | *31-35* | *7* |
| Syt1$^{+/+}$ Syt1$^{F349A}$ | *29-35* | *7* |

**Supplementary Figure 2b anti-Syt1 Ab**

| *Condition 1* | *Condition 2* | *P value* | *Statistical test* |
|---|---|---|---|
| Syt1$^{+/+}$ Control | Syt1$^{+/+}$ Syt1$^{WT}$ | P = <0.001 | Mann-Whitney U test |
| Syt1$^{+/+}$ Control | Syt1$^{+/+}$ Syt1$^{F349A}$ | P = <0.001 | Mann-Whitney U test |
| Syt1$^{+/+}$ Syt1$^{WT}$ | Syt1$^{+/+}$ Syt1$^{F349A}$ | P = 0.231 | Mann-Whitney U test |

**Supplementary Figure 2b anti-Myc Ab**

| *Condition 1* | *Condition 2* | *P value* | *Statistical test* |
|---|---|---|---|
| Syt1$^{+/+}$ Control | Syt1$^{+/+}$ Syt1$^{WT}$ | P = <0.001 | Mann-Whitney U test |
| Syt1$^{+/+}$ Control | Syt1$^{+/+}$ Syt1$^{F349A}$ | P = <0.001 | Mann-Whitney U test |
| Syt1$^{+/+}$ Syt1$^{WT}$ | Syt1$^{+/+}$ Syt1$^{F349A}$ | P = 0.851 | Mann-Whitney U test |

**Supplementary Figure 2c anti-Syt1 Ab**

| Condition 1 | Condition 2 | P value | Statistical test |
|---|---|---|---|
| $Syt1^{+/+}$ Control | $Syt1^{+/+}$ $Syt1^{WT}$ | P = 0.570 | Mann-Whitney U test |
| $Syt1^{+/+}$ Control | $Syt1^{+/+}$ $Syt1^{F349A}$ | P = 0.730 | Mann-Whitney U test |
| $Syt1^{+/+}$ $Syt1^{WT}$ | $Syt1^{+/+}$ $Syt1^{F349A}$ | P = 0.679 | Mann-Whitney U test |

**Supplementary Figure 2c anti-Myc Ab**

| Condition 1 | Condition 2 | P value | Statistical test |
|---|---|---|---|
| $Syt1^{+/+}$ Control | $Syt1^{+/+}$ $Syt1^{WT}$ | P = <0.001 | Mann-Whitney U test |
| $Syt1^{+/+}$ Control | $Syt1^{+/+}$ $Syt1^{F349A}$ | P = <0.001 | Mann-Whitney U test |
| $Syt1^{+/+}$ $Syt1^{WT}$ | $Syt1^{+/+}$ $Syt1^{F349A}$ | P = 0.712 | Mann-Whitney U test |

### <u>Supplementary Figure 3</u>

**Supplementary Figure 3b - Western Blot analysis**

| Condition | Number of cells | Number of preparations |
|---|---|---|
| $Syt1^{+/+}$ $Syt1^{WT}$ | | 3 |
| $Syt1^{+/+}$ $Syt1^{F349A}$ | | 4 |
| $Syt1^{+/+}$ $Syt1^{WT}$ Syt7 KD | | 3 |
| $Syt1^{+/+}$ $Syt1^{F349A}$ Syt7 KD | | 4 |

**Supplementary Figure 3b anti-Syt7 Ab**

| Condition 1 | Condition 2 | P value | Statistical test |
|---|---|---|---|
| $Syt1^{+/+}$ $Syt1^{WT}$ | $Syt1^{+/+}$ $Syt1^{WT}$ Syt7 KD | P = 0.007 | Student t test |
| $Syt1^{+/+}$ $Syt1^{F349A}$ | $Syt1^{+/+}$ $Syt1^{F349A}$ Syt7 KD | P = 0.008 | Student t test |

**Supplementary Figure 3b anti-Syt1 Ab**

| Condition 1 | Condition 2 | P value | Statistical test |
|---|---|---|---|
| $Syt1^{+/+}$ $Syt1^{WT}$ | $Syt1^{+/+}$ $Syt1^{WT}$ Syt7 KD | P = 0.846 | Student t test |
| $Syt1^{+/+}$ $Syt1^{F349A}$ | $Syt1^{+/+}$ $Syt1^{F349A}$ Syt7 KD | P = 0.407 | Student t test |

**Supplementary Figure 3d - Immunostaining**

| Condition | Number of cells | Number of preparations |
|---|---|---|

| | | |
|---|---|---|
| $Syt1^{+/+}$ $Syt1^{WT}$ | *15* | *3* |
| $Syt1^{+/+}$ $Syt1^{F349A}$ | *12* | *3* |
| $Syt1^{+/+}$ $Syt1^{WT}$ Syt7 KD | *12* | *3* |
| $Syt1^{+/+}$ $Syt1^{F349A}$ Syt7 KD | *12* | *3* |

**Supplementary Figure 3d  anti-Myc Ab**

| *Condition 1* | *Condition 2* | *P value* | *Statistical test* |
|---|---|---|---|
| $Syt1^{+/+}$ $Syt1^{WT}$ | $Syt1^{+/+}$ $Syt1^{WT}$ Syt7 KD | P = 0.903 | Mann-Whitney U test |
| $Syt1^{+/+}$ $Syt1^{F349A}$ | $Syt1^{+/+}$ $Syt1^{F349A}$ Syt7 KD | P = 0.931 | Mann-Whitney U test |

**Supplementary Figure 3d anti-Syt7 Ab**

| *Condition 1* | *Condition 2* | *P value* | *Statistical test* |
|---|---|---|---|
| $Syt1^{+/+}$ $Syt1^{WT}$ | $Syt1^{+/+}$ $Syt1^{WT}$ Syt7 KD | P = <0.001 | Mann-Whitney U test |
| $Syt1^{+/+}$ $Syt1^{F349A}$ | $Syt1^{+/+}$ $Syt1^{F349A}$ Syt7 KD | P = <0.001 | Mann-Whitney U test |
